# Quantifying the distribution of feature values over data represented in arbitrary dimensional spaces

**DOI:** 10.1101/2022.11.23.517657

**Authors:** Enrique R. Sebastian, Julio Esparza, Liset M de la Prida

## Abstract

**Background:** Identifying the structured distribution (or lack thereof) of a given feature over a point cloud is a general research question. In the neuroscience field, this problem arises while investigating representations over neural manifolds (e.g., spatial coding), in the analysis of neurophysiological signals (e.g., auditory coding) or in anatomical image segmentation.

**New method:** We introduce the Structure Index (SI) as a graph-based topological metric to quantify the distribution of feature values projected over data in arbitrary D-dimensional spaces (neurons, time stamps, pixels). The SI is defined from the overlapping distribution of data points sharing similar feature values in a given neighborhood.

**Results:** Using model data clouds we show how the SI provides quantification of the degree of local versus global organization of feature distribution. SI can be applied to both scalar and vectorial features permitting quantification of the relative contribution of related variables. When applied to experimental studies of head-direction cells, it is able to retrieve consistent feature structure from both the high- and low-dimensional representations. Finally, we provide two general-purpose examples (sound and image categorization), to illustrate the potential application to arbitrary dimensional spaces.

**Comparison with existing methods:** Most methods for quantifying structure depend on cluster analysis, which are suboptimal for continuous features and non-discrete data clouds. SI unbiasedly quantifies structure from continuous data in any dimensional space.

**Conclusions:** The method provides versatile applications in the neuroscience and data science fields

**Highlights:**

- The Structure Index is a graph-based topological metric
- It quantifies the distribution of feature values in arbitrary dimensional spaces
- It can be applied to both scalar and vectorial features
- When applied to the head-direction neural system, it extracts concordant information from high- and low-dimensional representations
- It can be extended to sound and image categorization, expanding the range of applications

## 1. Introduction

Identifying and quantifying if and how a given feature is structured along a data cloud is a challenging problem in many fields of science. For instance, in the neurosciences the temporal evolution of neuronal activity can be pictured as a data cloud on a high-dimensional space, whose axes are determined by the number of simultaneously recorded cells or recorded channels (Churchland et al., 2012). Under certain conditions, the high-dimensional activity can be embedded into 2D or 3D subspaces, where external and internal variables are reliably visualized (Cunningham and Yu, 2014; Nieh et al., 2021).Examples include neural manifolds underlying simple motor tasks (Gallego et al., 2017) and the internal head-direction and grid-cell representational systems (Chaudhuri et al., 2019; Gardner et al., 2022). In this context, understanding how a given feature is topologically organized over the manifold sheds light into the representational capacity of the system under study.

Other applications include analysis of multidimensional data that reflect temporal samples, such as the auditory coding of the spectro-temporal features of natural sounds (Gervain and Geffen, 2019), image segmentation of multidimensional pixels (Ternes et al., 2022) or transcriptomic data (Zeisel et al., 2015). In most cases, evaluating the unknown distribution of a feature over data samples (e.g., the motor reach, spatial and speech representations, histological categories in an image or neurodevelopmental profiles across cell-type clusters) relies on the visual inspection of the reduced embedding. Whether the very same feature had structure in the original high-dimensional space typically remains unclear.

Solving this general-purpose problem can provide solutions for an ample set of scientific applications. Having the ability to quantify the feature structure in any arbitrary space (i.e., that defined by cells, temporal samples, or pixels) may boost applications across fields. Here, we use the term structure in a loose sense. That is, we say that a variable or feature is structured along data if it follows some type of non-random distribution in the D-dimensional representational space.

Most methods for structure quantification depend on clustering analysis. However, when data points do not aggregate in groups or the features do not take discrete or nominal values, the resulting clusters are not directly interpretable. This renders these methods suboptimal to many real-world problems where continuous variables and point distributions are the norm. To overcome this limitation, some studies resort to techniques that depend on linear correlation metrics, posing limitations for the analysis of more realistic convoluted distributions (Enns, 2011). Alternative approaches based on decoders tacitly assume that if a given variable can be decoded from the data cloud, then it must follow some structure. However, this kind of strategies are highly dependent on the model used, as well as on the intrinsic dimensionality of the data, being vulnerable to overfitting as sparsity increases with dimensionality. Crucially, all these approaches provide poor insights about the local versus global structure of feature representations. It is therefore important to develop a method that (i) can be applied to non-linear distributions, (ii) generalizes to continuous features, and (iii) is applicable to arbitrarily high-dimensional spaces.

In this paper, we introduce the Structure Index (SI) as a new metric specifically aimed at quantifying how a given feature is topologically organized along an arbitrary data cloud. We first demonstrate the principles of our approach with simple model examples and illustrate how the method can be tuned to quantify the degree of the local/global organization of feature distribution, as well as its robustness along a broad range of data characteristics. We show how the SI can be equally applied to vectorial features, in which more than one variable can be considered. Next, we apply the SI to neural data from experimental studies of head-direction cells, showing how it can retrieve representation of different features, which are quantified beyond visual inspection of the neural manifold. Finally, we provide two additional general-purpose examples (sound and image categorization), to illustrate the universal application to topological data analysis across fields.

## 2. Material and Methods

### 2.1. Definition of the Structure Index (SI)

The SI aims at quantifying the amount of structure present at the distribution of a given feature over a point cloud in an arbitrary D-dimensional space. For instance, feature values can be distributed in a 2D cloud along a gradient (Fig.1A), or randomly (Fig.1E). Identifying such structure without the need for visualization is a major problem in many applications, especially for high-dimensional spaces. In the neuroscience field, this problem arises for instance when relating the distribution of neuronal activity to external behavioral variables projected over the neural manifold.

**Fig. 1.**
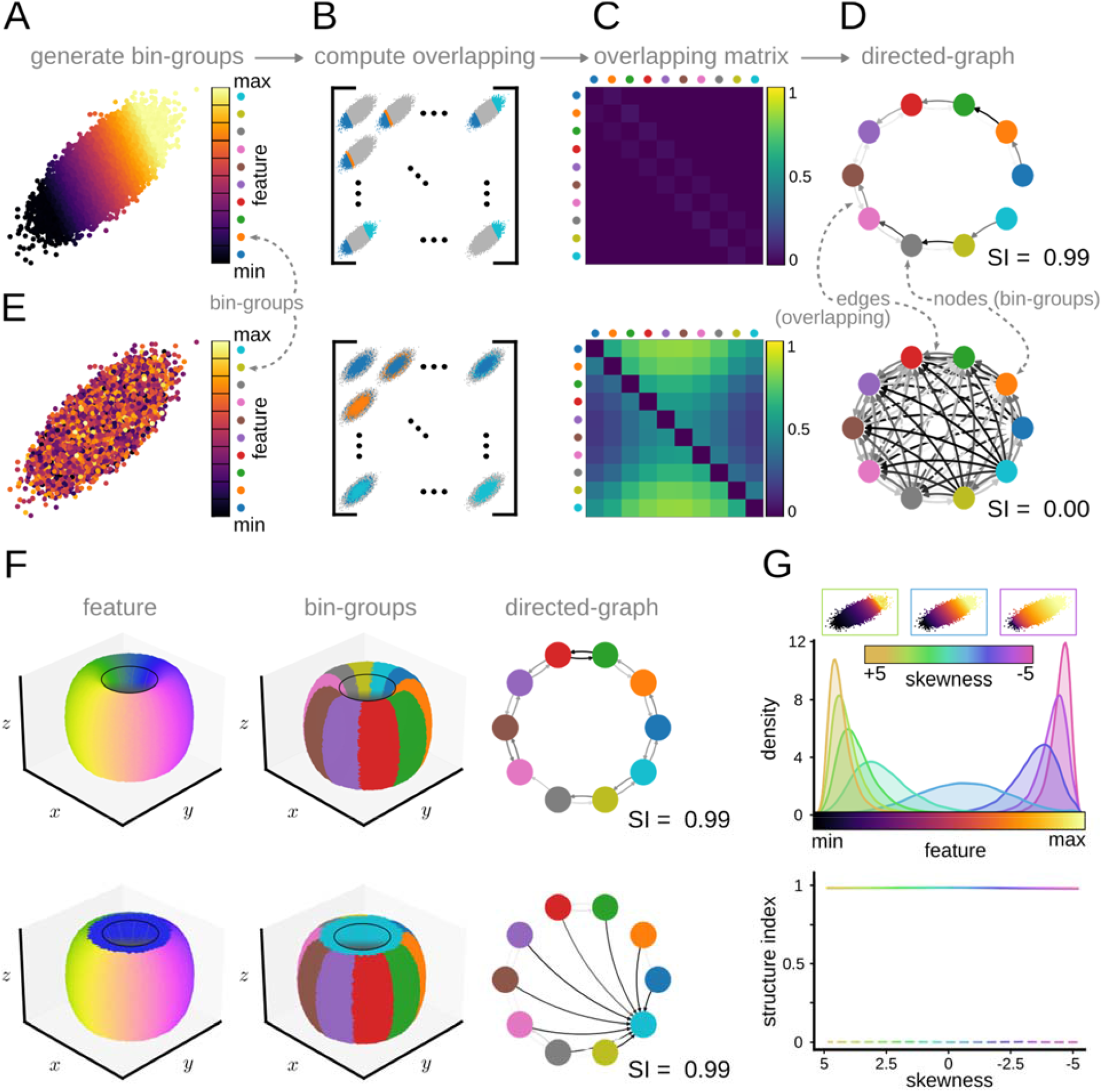
Illustration of the concepts behind the definition of the Structure Index (SI). **A**, Feature gradient distribution in a 2D-ellipsoid data cloud. Each point in the data cloud is assigned to a group associated with a feature bin value (bin-group). **B, C**, Next, the overlapping matrix between bin-groups is computed according to equation 1. **D**, The overlapping matrix represents a weighted directed graph between bin-groups, where structure (overlapping, clustering, etc..) can be quantified using the SI from 0 (random, equivalent to full overlapping) to 1 (maximal separation, equivalent to zero overlapping between bins). **E**, The case of a feature randomly distributed over a 2D data cloud. **F**, Different feature distribution yielding the same SI but different weighted directed graph. **G**, Lack of effect of the skewness of feature values on the SI.

To quantify feature distribution over a point cloud, we first divide the range of values in n-equal bins, and then assign each data point to a bin-group according to its feature value (Fig.1A). Note that features can be either categorical (i.e. they may take nominal values associated to different categories) or continuous (i.e. they may take values within a scalar range). In the case of a discrete feature, each bin-group may correspond to one of the possible discrete or nominal values the feature can take.

Next, we compute the overlapping between each pair of bin-groups in terms of the k-nearest neighbors (Fig.1B). Given two bin-groups, *u* and *v*, we define the overlapping score from *u* to *v* (*OS*_*u* →*v*_) as the ratio of k-nearest neighbors of all the points of *u* that belong to *v* in the point cloud space. That is,

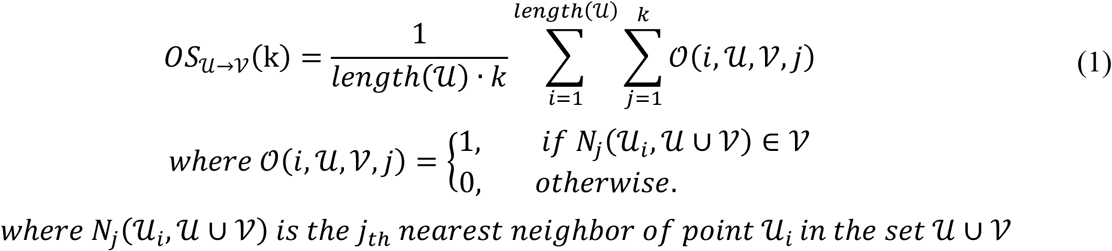

Note that the definition of nearest neighbors is determined by the distance metric used (i.e., Euclidean distance, geodesic distance, etc.). Computing the overlapping score for each pair of bin-groups (*u*_*a*_ and *v*_*b*_) yields an adjacency matrix (*ℳ*_*nxn*_) whose entry (*a, b*) equals 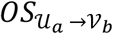 (Fig. 1C). *ℳ* can be thought as representing a weighted directed graph, where each node is a bin-group, and the edges represent the overlap (or connection) between them (Fig.1D).

Finally, we define the Structure Index as 1 minus the mean weighted degree of the nodes after scaling it:

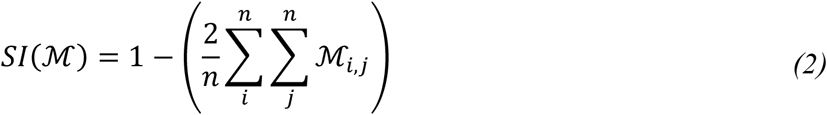

Under this definition, for a uniform random distribution the overlapping of any two nodes would be equal to 0.5 and therefore, the mean degree of the nodes of such distribution would also be 0.5. Thus, the Structure Index would take a value of 0 for a random distribution. In contrast, the mean degree of the nodes of a perfectly separated distribution would be 0 and thus, the SI would be 1. Therefore, the SI ranges between 0 (random feature distribution, fully connected graph) and 1 (maximally separated feature distribution, non-connected graph; Fig. 1D).

By definition, the SI is agnostic to the type of structure (e.g., gradient, patchy, etc.) since bin-groups do not need to follow any specific arrangement. Instead, the weighted directed graph provides additional insights. Fig. 1F shows the example of two different distributions with similar SI but different graphs. Of note, the skewness of feature values has little impact on SI, being robust for a wide range of statistical properties (Fig. 1G).

Note that this metric can be applied to any type of data represented in arbitrary D-dimensional spaces (cells, time series, pixels). Our approach is not in direct competition with the many methods that use cluster analysis or topological decoding. Rather, it generalizes at a class of distributions (i.e., continuous distributions) where clusters typically fail to apply. Our definition of SI and the equivalent graph makes this metric general enough to ease a range of application, which we will illustrate along the Results section.

### 2.2. Parameter dependence of SI on the neighborhood size

To compute the overlapping between each pair of bin-groups, the SI looks at the properties of the *k*-nearest neighbors of each point. For a low number of neighbors, the overlapping is computed in the close vicinity of each point, thus being biased towards the local distribution. As the number of neighbors increases, the SI tends to better account for the global structure. This dependence of the SI on the number of neighbors can be exploited to infer information about the local versus global organization of data features.

Figure 2 shows two different feature distributions over the same data cloud. In the local pattern, feature values replicate at the different regions of the data cloud and so bingroups reflect such organization (Fig. 2A). In contrast, in the global pattern, feature values follow a general trend (Fig. 2B). By evaluating the evolution of the SI as a function of the number of neighbors, the trade-off between local and global structure can be quantified.

**Fig. 2.**
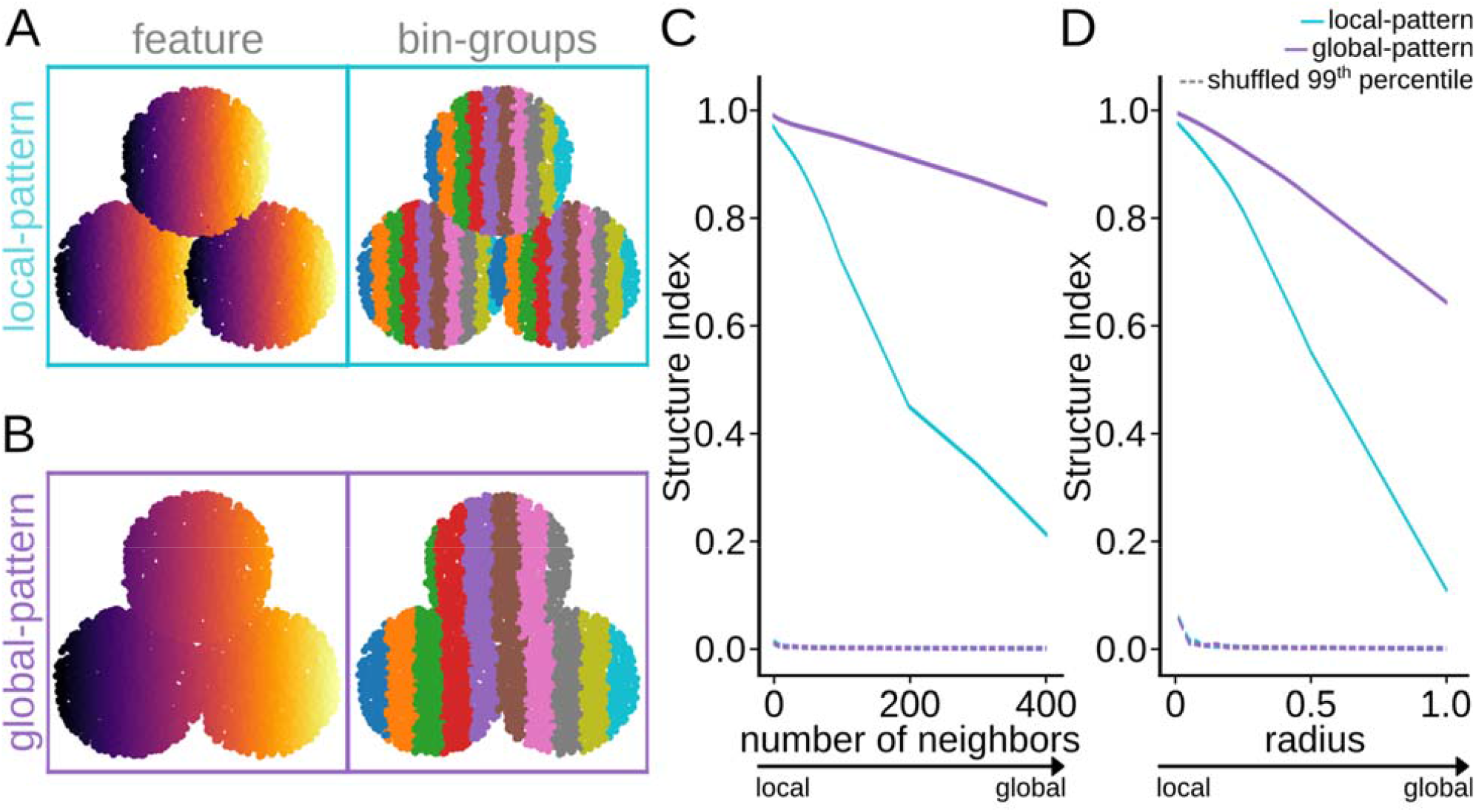
Parametric dependence of SI on the number of neighbors. **A**, A local pattern is simulated in 2D by projecting feature values differently along the data cloud (9000 points). Note local structure between bin-groups. **B**, The same 2D data cloud exhibiting a global distribution of feature values. **C**, Dependency of the SI values as a function of the number of neighbors can help to identify the local versus the global distribution trends. Data is tested against shuffled distribution of feature values (99^th^ percentile). **D**, Same as in C, but as a function of the radius.

For the local pattern, overlapping between bin-groups increases as the number of neighbors increase, and thus the SI sharply decreases. On the contrary, for the global pattern, the overlapping is less sensitive to the number of neighbors, and therefore the SI decreases smoothly (Fig. 2C). As expected, the SI of the shuffled distribution equals 0 independently on the number of neighbors. Thus, by tuning the number of neighbors, one can effectively change the sensitivity of SI to better detect local or global structures.

For data clouds with highly uneven density, the SI presents the option of setting a radius size (*r*) instead of the number of neighbors. In such a case, the neighbors of a point are set to be all points that fall within a given distance *r*. That is, equation (1) becomes:

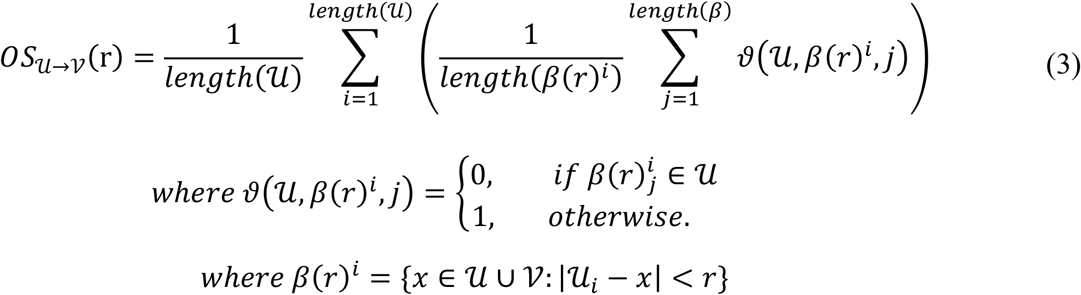

In such cases, the radius still helps to control for the trade-off between local and global structure, with smaller values making the SI more sensitive to local, and larger values being more sensitive to global structure (Fig. 2D).

### 2.3. Datasets

In this study, we used different datasets to evaluate SI performance. For the parameter study, we created objects (2D-ellipsoids, balls and spheres) using the corresponding mathematical equations. For the object lamp, we used the model from the ModelNet40 dataset, which is publicly available at https://github.com/antao97/PointCloudDatasets. Different feature value distributions were created over these objects and used to evaluate SI performance. By default all objects were created with 40,000 data points, except otherwise reported.

To study neural manifold representations, a publicly available head-direction dataset was used (http://crcns.org/data-sets/thalamus/th-1; doi:10.6080/K0G15XS1) (Peyrache et al., 2015). We chose this dataset because the neural manifold organization of head direction angles was recently validated (Chaudhuri et al., 2019), excluding any confounding in the ability of the SI to extract structure. Moreover, as we will show in the Results section, using these data allowed us to illustrate the capacity of the SI to quantify structure in the original space, which was not tested in the aforementioned reference due to lack of computationally efficient available methods. We used all data available to build the 3D neural manifold as reported in (Chaudhuri et al., 2019) using Isomap (Tenenbaum et al., 2000). We also built the representations in the original space using single cell data (one axis per cell), yielding different high-dimensional spaces per mouse (n=6; mouse12 - 120806: 37 cells; mouse17-130130: 29 cells; mouse20-130520: 11 cells; mouse24-131216: 10 cells; mouse25-140130: 10 cells; mouse28-140313: 22 cells). Information about brain states was used to separate the neural manifolds from awake and sleeping periods, with sleep classified as Slow-Wave Sleep (SWS) or Rapid Eyes Movement (REM) sleep (Chaudhuri et al., 2019).

To evaluate the application of SI to temporal data, we opted to use musical notes given their similarities with electroencephalographic waveforms (Baier et al., 2007). An additional advantage is that musical notes are directly interpretable, allowing us to focus in evaluating SI performance. We chose the NSynth dataset (Engel et al., 2017), which contains over 300,000 musical notes produced by around 1000 different acoustic, electronic or synthetic instruments, including the human voice. This dataset is available on the TensorFlow Magenta project at https://magenta.tensorflow.org/datasets/nsynth. Different features (source, instrument family, pitch and velocity) characterize each musical note. Each note consists in 4 seconds of monophonic 16 kHz audio snippets at five different velocities. For analysis, we downsampled the original snippets to 1.2 kHz resulting in 4800 time-stamps, which were used to build the high-dimensional space (one point per note). To comply with the Nyquist–Shannon sampling theorem, audio snippets with an associated pitch higher than 73 MIDI (equivalent to 554 Hz) were discarded. Binary features were not included in the analysis. For statistical testing, the dataset was divided in 5 equivalent batches, which were analyzed both in the original and the 3D-reduced space using Uniform Manifold Approximation and Projection (UMAP) (McInnes et al., 2018).

Finally, to provide examples of image analysis using SI we resorted to the bird species problem, given its application in fine-grained image recognition. Similar as above, this allowed us to focus in evaluating the performance of SI without requiring any particular interpretation. To this purpose, we used SI the 100-bird species dataset created by Gerald Piosenka, which is hosted on the Kaggle platform (https://www.kaggle.com/datasets/gpiosenka/100-bird-species; date of download July 28^th^, 2022). The dataset consists of more than 70000 RGB images of 450 bird species. Images are 224 × 224 pixels x 3 color (jpg format) annotated by species name. For analysis, we downsampled images to 56×56 pixels x 3 colors, resulting in a 9408-dimensional space, where each axis is the value of a pixel (one point per image). To expand the number of features associated to each image, we performed an automated data scraping from Wikipedia, so that for each bird species we also extracted information about geographical distribution (continents), as well as the scientific order and family, using the Python library ‘Wikipedia’ (https://pypi.org/project/wikipedia/). The dataset was divided in 2 equivalent batches.

Statistical analysis of different instances of each dataset was performed using one-or two-way ANOVAs followed by Student t-tests or equivalent. Spearman correlation was used to evaluate relationship between variables, which were fitted by exponential curves.

### 2.4. Computational resources

All simulations and analysis were performed in Python 3.8.13 using personal computer workstations (Intel Xeon CPU E5-2620 v4 @ 2.10GHz processor with 16 cores, 64GB RAM memory, GeForce GTX 1080 Ti GPU with 11GB memory and 0.355 TFlops for double precision). Whenever required, the supercomputer cluster Artemisa (https://artemisa.ific.uv.es/web/content/nvidia-tesla-volta-v100-sxm2) was used to accelerate calculation and parametric analysis (NeuroDIM Project).

### 2.5. Data and code availability

All data used in this study is publicly available (see section 2.3). Code is deposited at https://github.com/PridaLab/structure_index.

## 3. Results

### 3.1. SI quantifies the topological distribution of scalar feature values

Before applying our method to the study of neural data, we used toy model data to illustrate its performance and robustness to a wide range of point cloud characteristics. We generated 3 independent toy-models, including a 2D linear gradient (40,000 points), a 3D solid ball (40,000 points) where the feature was distributed along the radius, and a 3D lamp (32,000 points) whose feature varies in terms of the three axes (Fig. 3A).

**Fig. 3.**
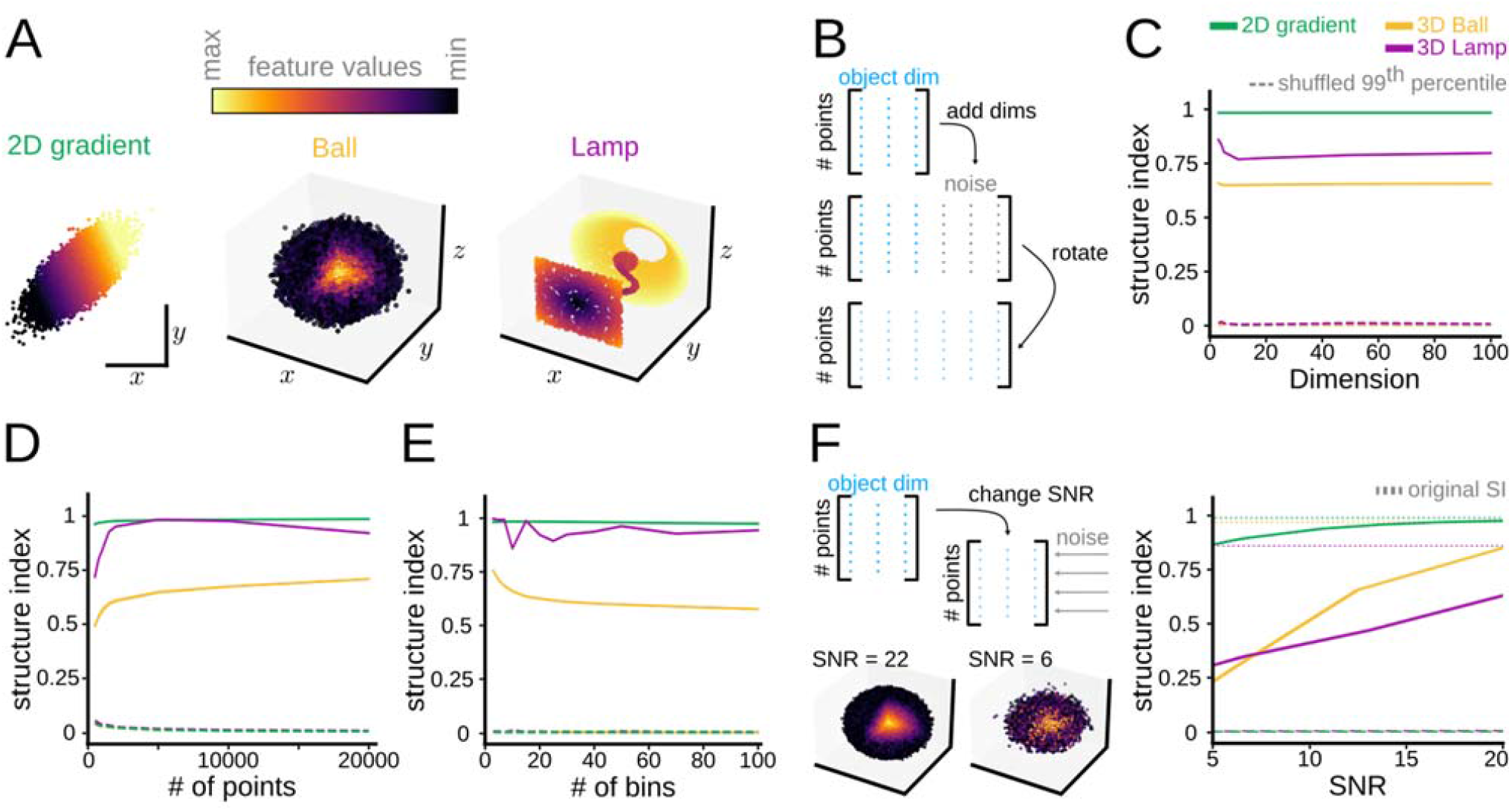
Robustness of SI under a wide range of data-cloud characteristics. **A**, Three toy models used to evaluate performance of SI (40,000 points for the gradient and the ball; 32,000 points for the lamp). **B**, Objects in A were embedded into spaces of increasing dimensionality by adding noise and then rotating. **C**, Dependence of the SI on the embedded dimensionality for the three toy models. **D**, Effects of the number of points in the data cloud as examined in a 2D space. **E**, Effects of the number of bin-groups on the SI for the three toy models. **F**, Effect of different levels of signal-to-noise ratio on SI.

To test for the stability of the method, we began by expanding these models in an increasing number of dimensions while adding white noise and then rotating the object in the extended space (Fig. 3B). By doing so, we maintained the intrinsic dimension of the object but spread the information along all dimensions. The SI showed a consistent response while increasing dimensionality (Fig. 3C). Importantly, the SI performed smoothly for a wide range of points in the cloud when examined in 2 dimensions (Fig. 3D).

In terms of the number of bin-groups used when computing the SI, there are two potential cases. For discrete or nominal features, the number of bin-groups is determined by the unique values the feature can take, so that there is a bin-group per discrete value. When dealing with continuous feature values, the number of bin-groups becomes a heuristic choice, which can be informed by statistical analysis. It should be large enough so that the continuity of the feature values is fully captured, but small enough so that there is a reasonable number of points assigned to each bin-group. While the SI performs consistently for a range of bin-groups (Fig. 3E), the topological characteristics of the data cloud may have different impacts that should be examined for each application.

Finally, we studied the sensitivity of the SI to different levels of noise in terms of the Signal to Noise Ratio (SNR), as defined in (Zeng et al., 2019). To this purpose, we introduced Gaussian noise across all existing dimensions (Fig.3F, left). While noise has effect in structure, the SI was able to capture the trends even when introducing high levels of noise into the point clouds (Fig. 3F, right). This renders the SI suitable for testing a wide variety of experimental data sets.

### 3.2. Evaluating the structure distribution of vectorial features

The definition of bin-groups used in the SI can be extended to vectorial features which integrate values from several characteristics. For example, a feature vector (*A, B*) can be created from two scalar features, *A* and *B*, taking values along a continuous scale. In such case, bin-groups can be defined by the upper and lower bound for both *A* and *B*. Thus, a point (*p*) in the cloud will fall within the bin-group *u* if and only if both entries of the associated feature vector fall within the common range.

To illustrate the case, we generated a point cloud sampled from a sphere of unitary radius using two angles *θ, φ*, with added Gaussian noise in 3D. Mathematically, the *x*-coordinate of a sphere is defined by the cosine of *θ*, while the *y*- and *z*-coordinates follow trigonometric relationships between both *θ, φ* (Fig. 4A). Thus, a feature defined by *θ* and *φ* independently will distribute differently along the sphere than a vectorial one integrating both angles (Fig. 4A). By definition, the structure of each angle separately should be lower than the vectorial angle (*θ, φ*). Moreover, given that the *x*-coordinate is completely defined by *θ*, we would expect more structure for *θ* than for *φ*. Consistently, the SI behaved as expected, with the lowest SI value obtained for *φ*, then for *θ*, and the highest value for both angles as a vectorial feature (*θ, φ*) (Fig. 4B).

**Fig. 4.**
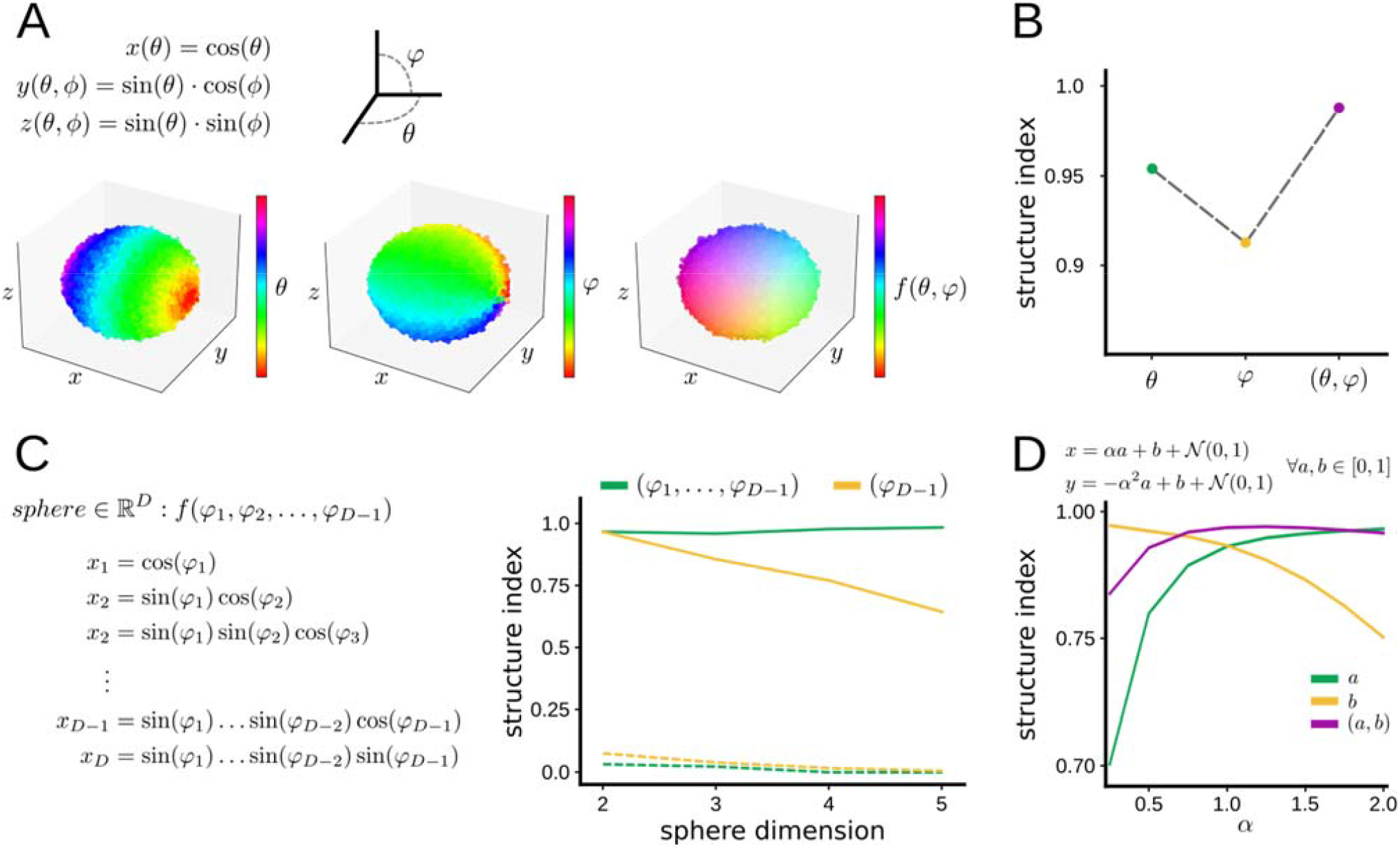
Evaluating structure of vectorial features. **A**, 3D sphere defined by trigonometric equations depending on angles *θ* and *φ* (40,000 points). Feature values can be defined for each angle independently, *θ* or *φ*, and for both together in a vectorial form (*θ, φ*). **B**, SI for each individual angle values and for the vectorial angle. **C**, A D-dimensional sphere is defined by trigonometric equations depending on *D*-1 angles (8^DxN^ points, with N=40,000 points to keep cloud density over D-dimensional spaces). The plot at right shows the dependence of SI on the sphere dimension, computed for the *D*-1 angle alone, and for all angles in vectorial form. Dashed lines indicate results from shuffled distribution values (99^th^ percentile). **D**, Behavior of SI for a feature defined in 2D according to the equation shown (20,000 points).

To evaluate the generalization of this behavior to vectorial features of any dimension, we generated point-clouds sampled from D-dimensional spheres according to the equation shown in Fig. 4C (left). For each point cloud in D-dimensional space, we computed the SI for both the *D* − 1 angle used to generate the sphere and all angles together as a feature vector (Fig. 4C, right). As predicted, the SI obtained when introducing all angles as a vector remained stable for all D-dimensional spheres. However, when only the *D* − 1 angle is considered, the SI declined as the dimensionality of the sphere increased. This reflects the fact that as the dimensions become larger, a lower percentage of coordinates depend on the *D*− 1 angle, and thus the position of a given point is less dependent on it.

This property of the SI can be exploited to examine the interdependence between distinct interrelated features. For instance, we created a 2D cloud where the position of each point depends on two features: *a, b* (Fig. 4D; see equation). While the impact of *b* in the position of the points was constant, the impact of *a* could be tuned by increasing or decreasing the parameter *α*. We proceeded by computing the SI of the scalar features *a, b*, and for the vector (*a, b*) using a range of *α* values (Fig. 4D). The maximum SI(*b*) was obtained for *α* equal to zero (as the position of the points was completely defined by *b*) and decreased consistently as *α* increased. In contrast, SI(*a*) increased with *α* as expected. Interestingly, SI(*a, b*) was lower than SI(*b*) for low *α* values (as the points are completely defined by *b*, the structure of (*a, b*) is lower than that of *b*). However, SI(*a, b*) rapidly increased with *α* reaching a plateau at maximum structure around 1 when both *a* and *b* equally contributed to the position of points.

These examples illustrate the capability of the SI to capture the structure of vectorial features, opening new avenues to study the relative impact and dependency between mathematically or experimentally related variables.

### 3.3. Application to neural manifold representations

Having established the main readouts expected from the SI metric, we sought to apply it to the study of neural manifolds. To illustrate the effectiveness of the approach, we chose a public dataset of extracellular recordings from multi-site silicon probes in the anterodorsal thalamic nucleus (ADn) of freely moving mice (Peyrache et al., 2015). This dataset has been recently used to demonstrate the intrinsic attractor manifold of the mammalian head-direction system (Chaudhuri et al., 2019), permitting direct testing of the ability of SI to extract feature structure.

In their study, Chaudhuri et al. showed that neural activity of N-simultaneously recorded ADn neurons of mice foraging in an open environment was constrained to a ring-shaped 3D manifold (Fig. 5A, right; n=6 mice), which they visualized in 3D using Isomap (Fig. 5B). Therefore, structure was implicitly expected at least in the low dimensional representation.

**Fig. 5.**
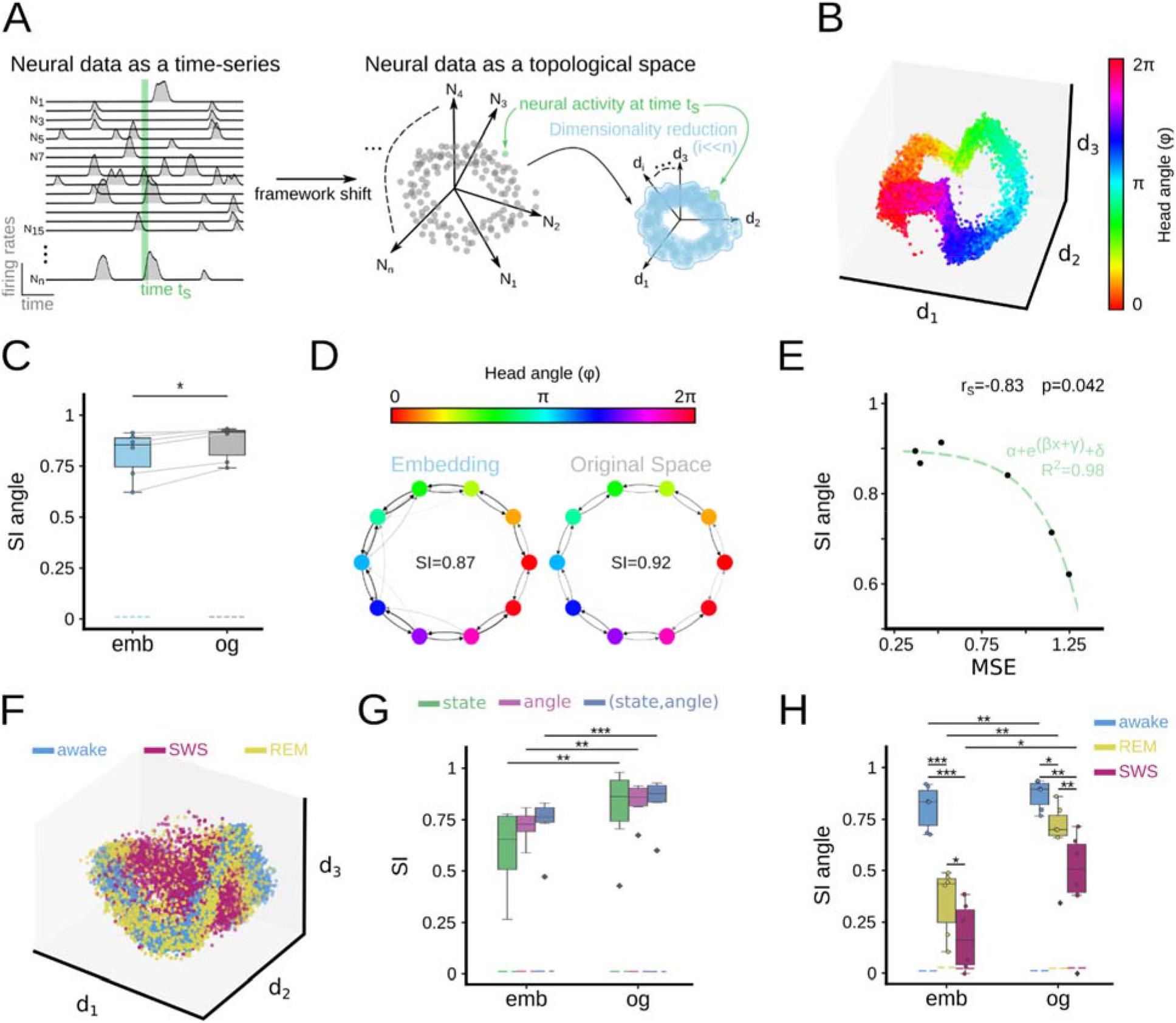
Using SI to evaluate neural representations. **A**, In the neural manifold framework, firing rates from N-neurons at a given time (t_s_) are represented in an N-dimensional Euclidean space. Activity is constrained in a subspace which can be retrieved using dimensionality reduction approaches (d-dimension). **B**, 3D neural manifold computed from the head direction system by Chaudhuri et al., with the head direction angle projected over the data cloud. **C**, SI of the head direction angle in the original (og) and 3D-embedded representations (emb). Dashed lines indicate results from shuffled distribution values (99^th^ percentile). **D**, Example of the weighted directed graphs from the same mouse (mouse12 - 120806) in the original and in the low-dimensional embedding. Note similar organization. **E**, Relationship between the SI and the mean square error (MSE) of the decoder trained by Chaudhuri et al. in the reduced space. Fitting curve parameters: *α*= -0.12; β=4.47; γ=-4.7 and Δ=0.9; tested significant at p<0.05. **F**, Head direction data plotted over the 3D embedding for awake, REM, and SWS states separately. **G**, SI for states and angles separately, and for both features together as expressed in a vectorial form. Results are shown for both the original (og) and the reduced space (emb). ANOVA effects for space (F(2,1)=8.2, p=0.007) but not for feature nor interaction. Post-hoc tests: *, p<0.05; **, p<0.01; ***, p<0.001. **H**, SI of the head direction angle for each state separately both in the original (og) and the reduced space (emb). ANOVA effects for state (F(2,1)=83.5, p<0.0001), space (F(2,1)=25.7, p<0.0001) and interaction. Post-hoc tests: *, p<0.05; **, p<0.01; ***, p<0.001.

When computing the SI of the head-direction angle over the neural manifold (3 neighbors), we obtained a high structure concordant with visual inspection of the embedding (Fig. 5C; blue). Importantly, in Chaudhuri et al. the analysis was mainly restricted to the 3D reduced space. Since the SI can be applied to an arbitrarily high-dimensional data cloud, we also evaluated the structure of the head-direction angle over the original N-dimensional neural space. Interestingly, the SI for all animals in the original space was slightly higher than in the low dimensional embedding (Fig. 5C; grey, paired sample t-test p=0.011). Visualization of individual weighted directed graphs from the high- and the low-dimensional representations confirm similar organization (Fig.5D).

In their original work, the author parametrized the manifold with splines of matching topology and used them to decode the represented latent variable (head-direction angle). We thus tested how the decoder performance (measured as the mean square error of predictions per mice) related to the head-direction information structured in the data. We found that the SI correlated with the decoder error (Spearman correlation -0.83, p=0.042), following an exponential decay relationship (R^2^=0.98; Fig.5E). That is, manifolds with lower decoding errors had higher head-direction structure as measured by SI.

Given the nature of the data, we wondered whether the head direction representation can be retrieved during REM as well as in SWS states (Senzai and Scanziani, 2022). To tackle this question, we resorted to the same dataset but used all neural data to compute the 3D manifold as reported in Chaudhuri et al. Indeed, when points of the manifold were colorcoded according to the state (awake, REM, nREM) we noted some stratification, which could be quantified using the SI (Fig. 5F). The SI returned structure for both the animal state and the head-direction angle, with higher values in the original than in the reduced space (Fig.5G; ANOVA effects for space, F(2,1)=8.2, p=0.007, but not for feature nor interaction). Interestingly, structure was higher when using a vectorial feature consisting on the state and the head-direction angle together, indicating that there may be some interdependency between them (Fig. 5G).

Finally, we computed the SI of the head-direction angle for each state separately (Fig. 5H). SI was maximal in awake conditions. In general, data represented in the original space provided more structure than in the manifold embedding (ANOVA effects for state (F(2,1)=83.5, p<0.0001), space (F(2,1)=25.7, p<0.0001) and interaction). Moreover, whereas REM and SWS yielded a low SI in the manifold, it was significantly higher in the original space, indicating that information was lost while reducing dimension. Thus, being able to evaluate neural activity in the original space using the SI might provide new insights into the representative capacity during multiple brain states.

### 3.4. Application to arbitrary D-dimensional spaces (temporal samples and images)

Finally, we applied the SI to two additional general-purpose examples: sound (temporal data) and image categorization (pixels), illustrating the usefulness of the SI metric for analysis of different types of data and across fields.

For temporal data, we resorted to musical notes given similarities with electro-encephalographic waveforms (Baier et al., 2007). In addition, using this dataset allowed us to focus in evaluating SI performance directly, given direct interpretability of musical notes. Data consisted on 4 seconds of musical notes of different pitch and velocity downsampled as 4800 time stamps. They were produced by different instruments (including the human voice) using acoustic, electronic or synthetic sources. Instruments are annotated as belonging to different families. Notes were represented in the 4800-dimensional space (Fig.S1A), and the SI was calculated both locally (using 3 neighbors) and globally (60 neighbors). In general, data showed a higher local than global structure (Fig.S1B; ANOVA effects F(3,1)=4.0, p<0.0001). We found that the pitch provided maximal structure, followed by the source and family, as confirmed by the weighted directed graph returned by the overlapping of instrument families (Fig.S1C; ANOVA effects for features F(3,1)=12.0, p<0.0001). Reducing data to 3D allowed for visualization of these trends (Fig.S1D), and provided similar SI figures as for the original space (Fig.S1E).

For image analysis, we chose using images of one hundreds bird species, typically exploited in fine-grained recognition problems. Data consist on RGB images (56×56×3 pixels) so that each image was represented as a point in a 9408-dimensional space (Fig.S2A). Birds were classified as belonging to different species, continent, scientific order and family. SI was maximal for bird species, followed by family and order (Fig.2SB). We noted that continents provided the lower structure, potentially reflecting migratory habits and/or species diversification. Visualization of images showing maximal and minimal overlapping values confirmed that the SI successfully captured the underlying structure of the data (Fig. S2C,D).

These examples illustrate how the SI method successful operates in arbitrary D-dimensional spaces, allowing for a range of multidisciplinary applications in neuroscience, as well as across other research fields.

## 4. Discussion

With the development of a graph-based topological metric (SI), we have enabled accurate quantification of the structure of feature distributions. The approach is not constrained by the dimensionality of the space and is robust to a wide range of data and feature characteristics. Importantly, the SI not only quantifies the “amount” of structure of scalar feature values represented over a point cloud, but it can also provide insights into the topological distribution of the feature by looking at the overlapping directed graph. Importantly, the SI is not limited to Euclidean spaces, as one can define the *k*-closest neighbors in terms of different distance metrics. For instance, the SI allows for the use of geodesic distance and cosine distance among others.

A common issue in current dimensionality reduction methods is being able to capture the global structure without deforming local relationships. Indeed, most dimensionality reduction methods have a parameter to control that tradeoff (e.g., the number of neighbors). Here, we demonstrated that the SI can be tuned to better detect local vs global structure by changing the number of neighbors (or equivalently the radius) used to compute overlapping between bin-groups. Thus, the SI can be used not only to quantify the structure in the original space, but also to evaluate the quality of the dimensionality reduction by looking at how much structure has been preserved both locally and globally while reducing from the high-to the low-dimensional representations.

As demonstrated above, the SI can be extended to vectorial features, expanding the range of applications. Note that a vectorial label can be created by grouping multiple scalar/categorical features, or by integrating several related variables. By doing so, the SI allows for the study of how different features interact with each other, allowing for a deeper understanding of how data structure is determined. This may ease data-driven discoveries of latent interaction between experimental features, which cannot be established a priori.

In this context, we have applied the SI to study the representation capability of the head-directional system (Peyrache et al., 2015). By using data from a previous study that demonstrated low dimensional representations of the head-direction angle (Chaudhuri et al., 2019), we have shown that the SI captures structure both in the lower dimensional manifold and in the original space. Moreover, by applying the metrics to awake, REM, and nREM states, we showed that the head-direction representation is preserved during REM, providing additional interpretation. This is consistent with recent data supporting mental replay of head-direction angles during REM sleep (Senzai and Scanziani, 2022).

The SI method can be applied to the study of temporal data expressed in high-dimensional spaces. By representing temporal events in the space built from the individual time stamps, electrophysiological signals can be analyzed with state space methods (Durbin and Koopman, 2001; Gervasoni et al., 2004; Reichinnek et al., 2010). Applying the SI to these representations may thus allow for new strategies for the analysis of the spectro-temporal organization of brain oscillations and/or perception (Valero et al., 2017; Lopes-Dos-Santos et al., 2018; Gervain and Geffen, 2019; Navas-Olive et al., 2020; Douchamps et al., 2022). Similarly, the SI permits image quantification and categorization in the service for fine-grained image recognition problems applicable to several research fields.

As topological and high-dimensional analysis become the norm in the neuroscience field, we expect that the SI will be a powerful tool to shed light into a wide range of questions. Here we have provided several examples, from high-dimensional geometrical analysis to sound and image categorization, expanding the applicability of the tool across fields.

## CRediT authorship contribution statement

ERS: Conceptualization, Methodology, Software, Validation, Formal analysis, Investigation, Writing. JE: Conceptualization, Methodology, Software, Validation, Formal analysis, Investigation, Writing. LMP: Conceptualization, Writing, Supervision Resources, Project Administration, Funding acquisition.

## Acknowledgments

This work is supported by a grant from Fundación La Caixa (LCF/PR/HR21/52410030; DeepCode) to LMP. JE received the support of a PhD fellowship from “la Caixa” Foundation (ID 100010434; LCF/BQ/DR22/11950026). Access to supercomputer cluster Artemisa (NeuroDIM) is co-funded by the European Union through the 2014-2020 FEDER Operative Programme of Comunitat Valenciana, project IDIFEDER/2018/048.

## Data availability

All data used in this study is publicly available (see section 2.3).

