## Supplementary figures for "Quantifying the distribution of feature values over data represented in arbitrary dimensional spaces"

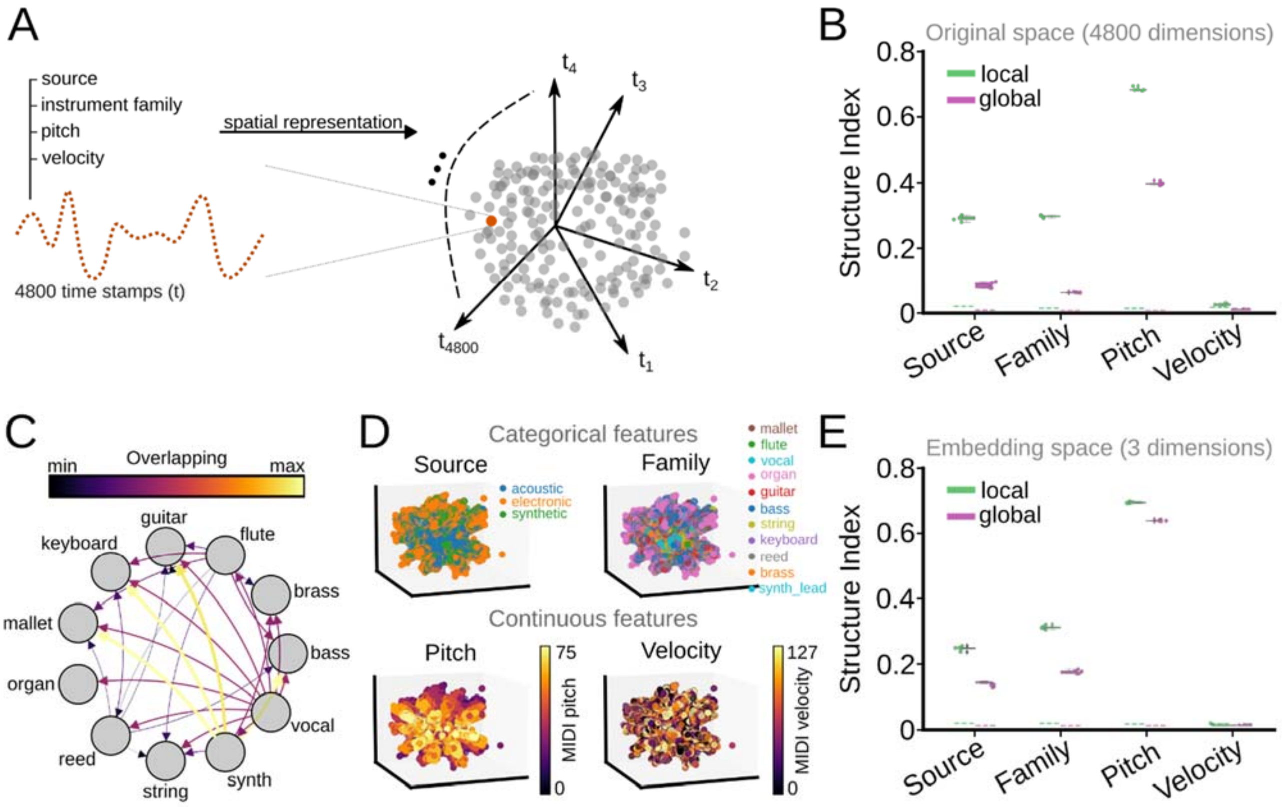


**Fig. S1. SI method applied to temporal data**. **A,** Data consists of musical notes from different instruments, which can be represented in a 4800-dimensional Euclidean space, where each axis is one timestamp. Notes of similar pitch are expected to lie closer in the high-dimensional space, with family instruments providing some additional structure. **B,** SI of the different features of the musical notes (source, instrument family, pitch, and velocity) both in a local (3 neighbors) and global (60 neighbors) region in the original space. Dashed lines represent 99$^{th}$ shuffled percentile. Global structure was in general lower as compared with local structure ($F(3,1)=4.0$, $p<0.0001$). Note higher structure for the pitch versus the source and instrument family ($F(3,1)=12.0$, $p<0.0001$. Individual data points represent results from 5 equivalent batches from the full dataset. **C,** Weighted directed graph returned by the overlapping of instrument family. Note nodes of similar instruments located closer in the graph. Note that direction and width of connecting edges give information on instrument similarity from different families. **D,** Projection of the different categorical and continuous features in a 3D embedding created with UMAP from the original 4800-dimensional space. Note larger structure for pitch than for source and family. **E,** SI of the different features of the musical notes in the 3D embedding both in a local (3 neighbors) and global (60 neighbors) vicinity. Dashed lines represent 99$^{th}$ shuffled percentile. Individual data points *represent results* from 5 equivalent batches from the full dataset.

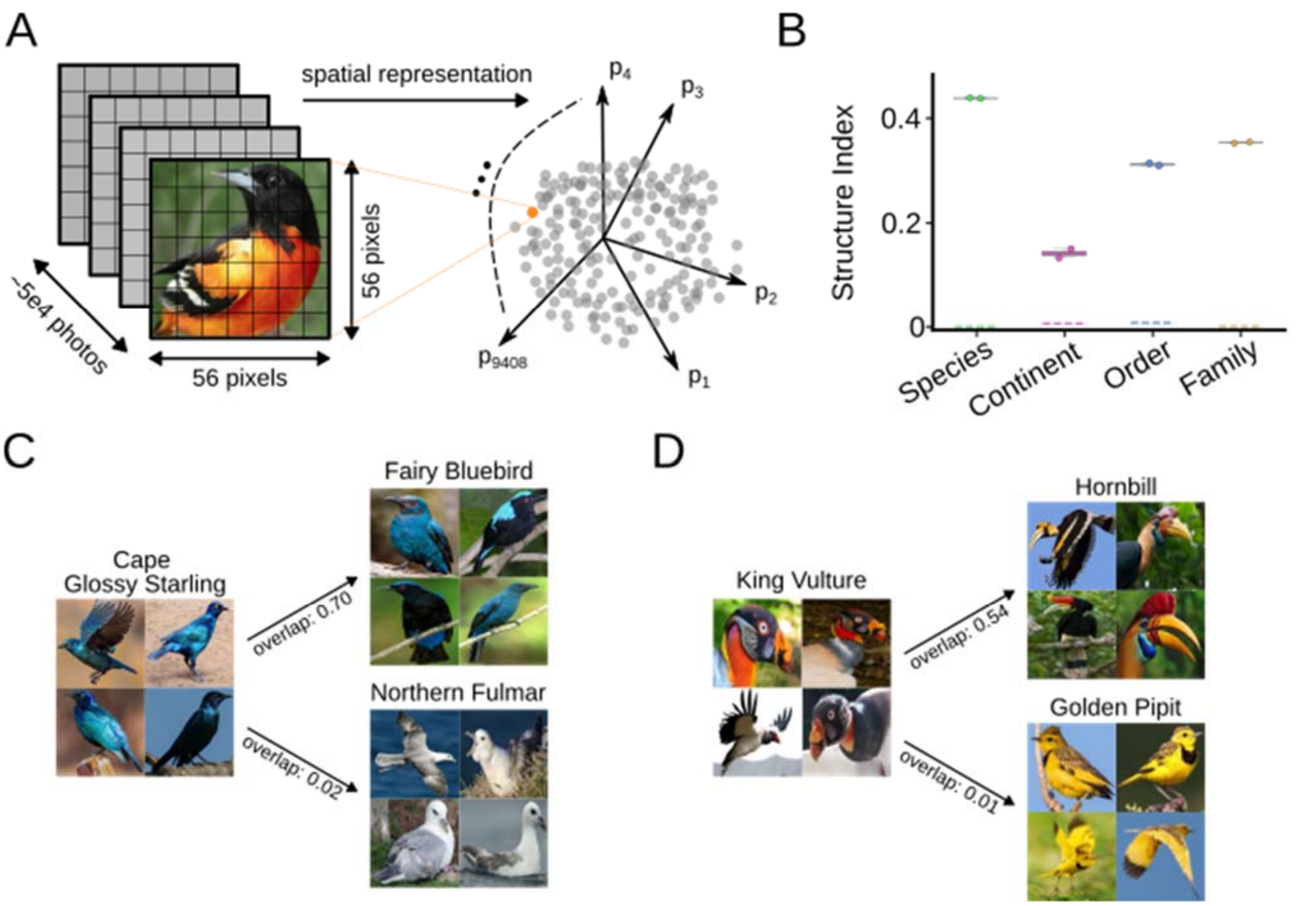


**Fig. S2. SI applied to image classification**. **A,** Data consists of annotated RGB images (56x56x3) of multiple bird species. Each image can be represented as a point in a 9408-dimensional Euclidean space where each axis is the value of a pixel. **B,** SI of the different features from each image, including the species, continent, scientific order and family. Dashed lines represent $99^{th}$ shuffled percentile. Individual data points represent results from 2 equivalent batches from the full dataset. **C,** Examples of species of birds showing maximal (Fairy Bluebird) and minimal overlap (Northern Fulmar) with Cape Glossy Starling. **E,** Examples of species showing maximal (Hornbill) and minimal overlap (Golden Pipit) with King Vulture.
